# MagicLamp: a web server and software toolkit for targeted gene annotation of microbial functions

**DOI:** 10.64898/2026.07.19.739457

**Authors:** A.I. Garber, I.A. Viney, N. Merino, G.A. Ramírez, M.J. Pavia, S. M. McAllister, S. Sadeghpour, A. Manna, M. Kreger, B. Wolk, E. Kim, J. Qu, C.R. Armbruster, I. Pérez-Rodríguez

## Abstract

Genome and metagenome annotation tools designed for large databases are ill-suited to the discovery of specialized, ecologically relevant microbial functions. MagicLamp (https://github.com/Arkadiy-Garber/MagicLamp) is a modular command-line software toolkit that performs targeted functional gene annotation searches using curated collections of hidden Markov models (HMMs), each representing discrete microbial metabolic processes. MagicLamp is also available as a web server: https://midauthorbio.com/#magiclamp. This targeted approach enables sensitive and specific annotation of genes involved in defined microbial processes, allowing MagicLamp to serve as a dedicated repository for the annotation of specialized microbial functions currently overlooked in other databases and software. The server accepts unannotated genome assemblies or GenBank-formatted annotations to perform HMM-based searches against curated model sets with reproducible, model-specific bit-score thresholds. Automated results are returned as tabular summaries and interactive HTML reports containing cross-genome/metagenome comparisons.

## Introduction

Functional gene annotation of microbial genomes and metagenomes typically depends on large, all-encompassing reference databases: KEGG (Kanehisa, 1997), COGs (Tatusov, 1997; Galperin et al., 2021), Pfam (Sonnhammer, 1997), and TIGRFAMs (Haft, 2003). While this breadth of gene detection is necessary for whole-genome characterization, the annotation signal for specialized and underrepresented microbial functions within these large databases is diluted. To balance the breadth of large databases with the specificity required to accurately detect discrete gene functions, we have developed an alternative targeted annotation approach based on curated HMM collections. Curated model sets, each constructed with a specific biological function in mind, allow the use of model-specific scoring thresholds calibrated to the biological function of interest, improving detection confidence in gene annotation assignments.

We present MagicLamp, a modular web server and software toolkit developed for the targeted annotation of genes relevant to microbial ecology, environmental microbiology, and/or biotechnology (**Table 1**).

**Table 1.**
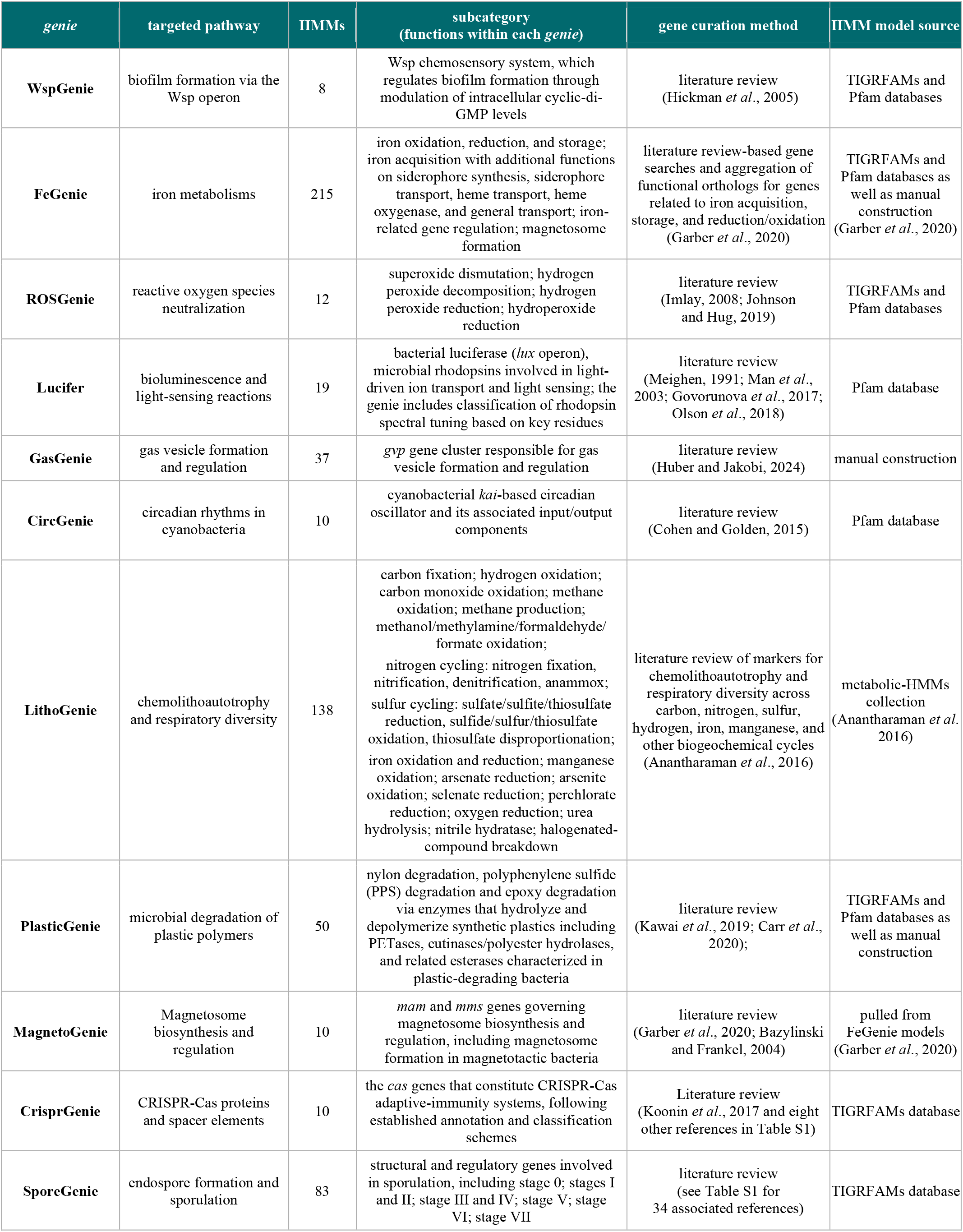
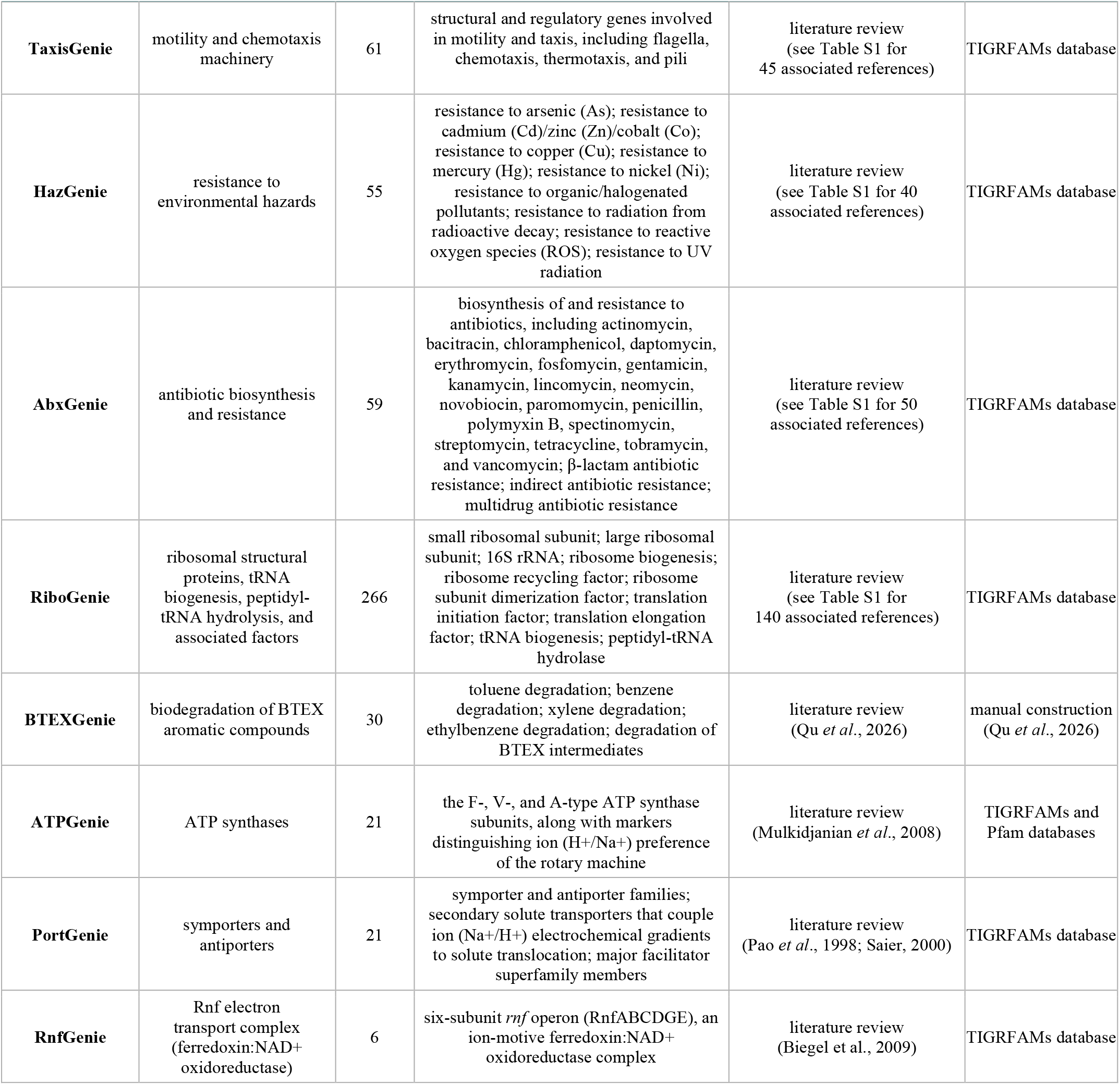
Summary of currently available *genie* modules (curated HMM collections) within MagicLamp.

| *genie* | targeted pathway | HMMs | subcategory<br>(functions within each *genie*) | gene curation method | HMM model source |
| --- | --- | --- | --- | --- | --- |
| **WspGenie** | biofilm formation via the Wsp operon | 8 | Wsp chemosensory system, which regulates biofilm formation through modulation of intracellular cyclic-di-GMP levels | literature review (Hickman *et al.*, 2005) | TIGRFAMs and Pfam databases |
| **FeGenie** | iron metabolisms | 215 | iron oxidation, reduction, and storage; iron acquisition with additional functions on siderophore synthesis, siderophore transport, heme transport, heme oxygenase, and general transport; iron-related gene regulation; magnetosome formation | literature review-based gene searches and aggregation of functional orthologs for genes related to iron acquisition, storage, and reduction/oxidation (Garber *et al.*, 2020) | TIGRFAMs and Pfam databases as well as manual construction (Garber *et al.*, 2020) |
| **ROSGenie** | reactive oxygen species neutralization | 12 | superoxide dismutation; hydrogen peroxide decomposition; hydrogen peroxide reduction; hydroperoxide reduction | literature review (Imlay, 2008; Johnson and Hug, 2019) | TIGRFAMs and Pfam databases |
| **Lucifer** | bioluminescence and light-sensing reactions | 19 | bacterial luciferase (*lux* operon), microbial rhodopsins involved in light-driven ion transport and light sensing; the *genie* includes classification of rhodopsin spectral tuning based on key residues | literature review (Meighen, 1991; Man *et al.*, 2003; Govorunova *et al.*, 2017; Olson *et al.*, 2018) | Pfam database |
| **GasGenie** | gas vesicle formation and regulation | 37 | *gvp* gene cluster responsible for gas vesicle formation and regulation | literature review (Huber and Jakobi, 2024) | manual construction |
| **CircGenie** | circadian rhythms in cyanobacteria | 10 | cyanobacterial *kai*-based circadian oscillator and its associated input/output components | literature review (Cohen and Golden, 2015) | Pfam database |
| **LithoGenie** | chemolithoautotrophy and respiratory diversity | 138 | carbon fixation; hydrogen oxidation; carbon monoxide oxidation; methane oxidation; methane production; methanol/methylamine/formaldehyde/formate oxidation;<br>nitrogen cycling: nitrogen fixation, nitrification, denitrification, anammox;<br>sulfur cycling: sulfate/sulfite/thiosulfate reduction, sulfide/sulfur/thiosulfate oxidation, thiosulfate disproportionation; iron oxidation and reduction; manganese oxidation; arsenate reduction; arsenite oxidation; selenate reduction; perchlorate reduction; oxygen reduction; urea hydrolysis; nitrile hydratase; halogenated-compound breakdown | literature review of markers for chemolithoautotrophy and respiratory diversity across carbon, nitrogen, sulfur, hydrogen, iron, manganese, and other biogeochemical cycles (Anantharaman *et al.*, 2016) | metabolic-HMMs collection (Anantharaman *et al.*, 2016) |
| **PlasticGenie** | microbial degradation of plastic polymers | 50 | nylon degradation, polyphenylene sulfide (PPS) degradation and epoxy degradation via enzymes that hydrolyze and depolymerize synthetic plastics including PETases, cutinases/polyester hydrolases, and related esterases characterized in plastic-degrading bacteria | literature review (Kawai *et al.*, 2019; Carr *et al.*, 2020); | TIGRFAMs and Pfam databases as well as manual construction |
| **MagnetoGenie** | Magnetosome biosynthesis and regulation | 10 | *mam* and *mms* genes governing magnetosome biosynthesis and regulation, including magnetosome formation in magnetotactic bacteria | literature review (Garber *et al.*, 2020; Bazylinski and Frankel, 2004) | pulled from FeGenie models (Garber *et al.*, 2020) |
| **CrisprGenie** | CRISPR-Cas proteins and spacer elements | 10 | the *cas* genes that constitute CRISPR-Cas adaptive-immunity systems, following established annotation and classification schemes | Literature review (Koonin *et al.*, 2017 and eight other references in Table S1) | TIGRFAMs database |
| **SporeGenie** | endospore formation and sporulation | 83 | structural and regulatory genes involved in sporulation, including stage 0; stages I and II; stage III and IV; stage V; stage VI; stage VII | literature review (see Table S1 for 34 associated references) | TIGRFAMs database |
| **TaxisGenie** | motility and chemotaxis machinery | 61 | structural and regulatory genes involved in motility and taxis, including flagella, chemotaxis, thermotaxis, and pili | literature review<br>(see Table S1 for 45 associated references) | TIGRFAMs database |
| **HazGenie** | resistance to environmental hazards | 55 | resistance to arsenic (As); resistance to cadmium (Cd)/zinc (Zn)/cobalt (Co); resistance to copper (Cu); resistance to mercury (Hg); resistance to nickel (Ni); resistance to organic/halogenated pollutants; resistance to radiation from radioactive decay; resistance to reactive oxygen species (ROS); resistance to UV radiation | literature review<br>(see Table S1 for 40 associated references) | TIGRFAMs database |
| **AbxGenie** | antibiotic biosynthesis and resistance | 59 | biosynthesis of and resistance to antibiotics, including actinomycin, bacitracin, chloramphenicol, daptomycin, erythromycin, fosfomycin, gentamicin, kanamycin, lincomycin, neomycin, novobiocin, paromomycin, penicillin, polymyxin B, spectinomycin, streptomycin, tetracycline, tobramycin, and vancomycin; β-lactam antibiotic resistance; indirect antibiotic resistance; multidrug antibiotic resistance | literature review<br>(see Table S1 for 50 associated references) | TIGRFAMs database |
| **RiboGenie** | ribosomal structural proteins, tRNA biogenesis, peptidyl-tRNA hydrolysis, and associated factors | 266 | small ribosomal subunit; large ribosomal subunit; 16S rRNA; ribosome biogenesis; ribosome recycling factor; ribosome subunit dimerization factor; translation initiation factor; translation elongation factor; tRNA biogenesis; peptidyl-tRNA hydrolase | literature review<br>(see Table S1 for 140 associated references) | TIGRFAMs database |
| **BTEXGenie** | biodegradation of BTEX aromatic compounds | 30 | toluene degradation; benzene degradation; xylene degradation; ethylbenzene degradation; degradation of BTEX intermediates | literature review<br>(Qu *et al.*, 2026) | manual construction<br>(Qu *et al.*, 2026) |
| **ATPGenie** | ATP synthases | 21 | the F-, V-, and A-type ATP synthase subunits, along with markers distinguishing ion (H<sup>+</sup>/Na<sup>+</sup>) preference of the rotary machine | literature review<br>(Mulikidjanian *et al.*, 2008) | TIGRFAMs and Pfam databases |
| **PortGenie** | symporters and antiporters | 21 | symporter and antiporter families; secondary solute transporters that couple ion (Na<sup>+</sup>/H<sup>+</sup>) electrochemical gradients to solute translocation; major facilitator superfamily members | literature review<br>(Pao *et al.*, 1998; Saier, 2000) | TIGRFAMs database |
| **RnfGenie** | Rnf electron transport complex (ferredoxin:NAD<sup>+</sup> oxidoreductase) | 6 | six-subunit *rnf* operon (RnfABCDGE), an ion-motive ferredoxin:NAD<sup>+</sup> oxidoreductase complex | literature review<br>(Biegel *et al.*, 2009) | TIGRFAMs database |

MagicLamp serves as a dedicated resource for targeted functional genome searches of microorganisms, where curated HMM collections, each representing a defined functional domain, are contained within individual *genies*. An advantage of this setup is that it is inherently extensible, facilitating the future incorporation of new functional gene modules needed for in-depth evaluation of difficult-to-assess microbial functions. Overall, MagicLamp offers a platform for comparative gene analysis and for generating hypotheses about gene homologs associated with targeted microbial capabilities.

### Software and Web Server Description

#### Architecture and Implementation

MagicLamp is implemented as a modular Python-based workflow that coordinates gene prediction, HMM searches, and automated organization and visualization of results (**Figure 1**). The software accepts unannotated genomic/metagenomic FASTA assemblies, which are annotated with Prodigal (Hyatt 2010), as well as GenBank-formatted annotation files (https://www.ncbi.nlm.nih.gov/genbank/samplerecord/).

**Figure 1.**
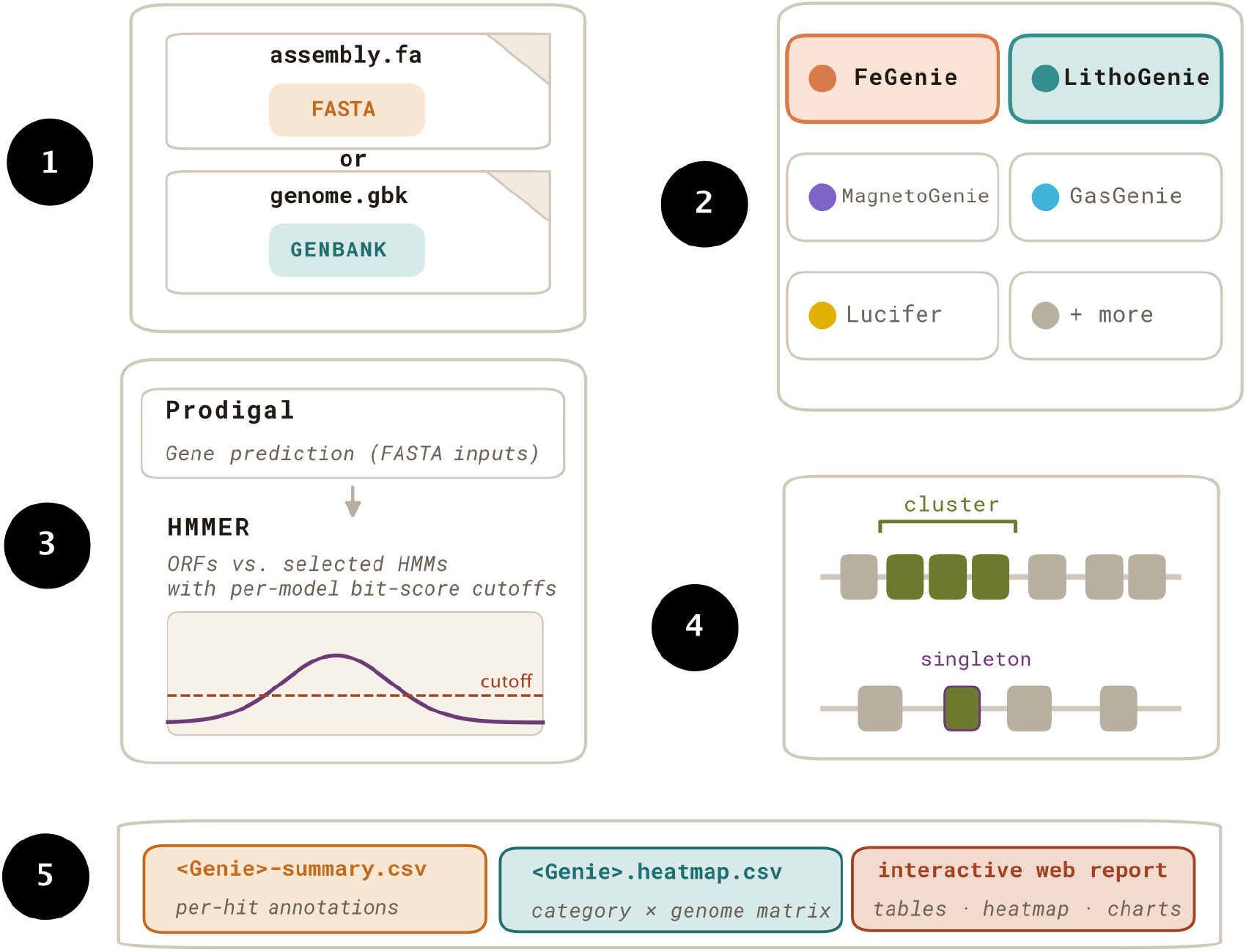
MagicLamp workflow: 1) User input can be raw genome/metagenome assemblies or annotated genomes in GenBank flat file format; 2) *Genie* selection (displayed in bold text) from a variety of gene pathways/categories; 3) HMM search is performed with HMMER after prediction of open reading frames (ORFs) with Prodigal, if raw assemblies in FASTA format are provided; 4) Gene proximity clustering is carried out to identify adjacent HMM hits. 5) Deliverables include two CSV files listing the results in matrix and long formats, as well as an interactive HTML report.

Functional annotation is performed using HMMER (Eddy 2008, 2009, 2011) with model-specific bit-score thresholds. The modular architecture of MagicLamp allows each *genie* module to be executed individually or in combination with other *genies* (**Figure 2A**), enabling users to tailor their analysis to specific research questions. Each *genie* represents a complete HMM collection for a target microbial function, along with associated metadata, bit score thresholds, and reference annotations. Following HMMER scanning, hits on the same contig whose open reading frames fall within a single gene neighborhood are assigned a shared cluster ID, allowing for inference of gene co-localization and possible operons (e.g., *luxCDABEG* cassettes, *mam* islands, or *gvp* operons). Importantly, many HMMs are not one-to-one with genes: a single HMM can map to multiple genes depending on biological context and function (for example, TIGR01494.1 matches *copB, zntA, copA, gph, kdpB*, and *cadA*). Because of the multiple functions generally associated with individual genes, a raw HMM hit alone is often not conclusive. By investigating the co-localization of genes along contigs, in addition to the co-occurrence of genes within a single genome or metagenome, ambiguous or multifunctional profiles can be interpreted in their proper genomic and functional context.

**Figure 2.**
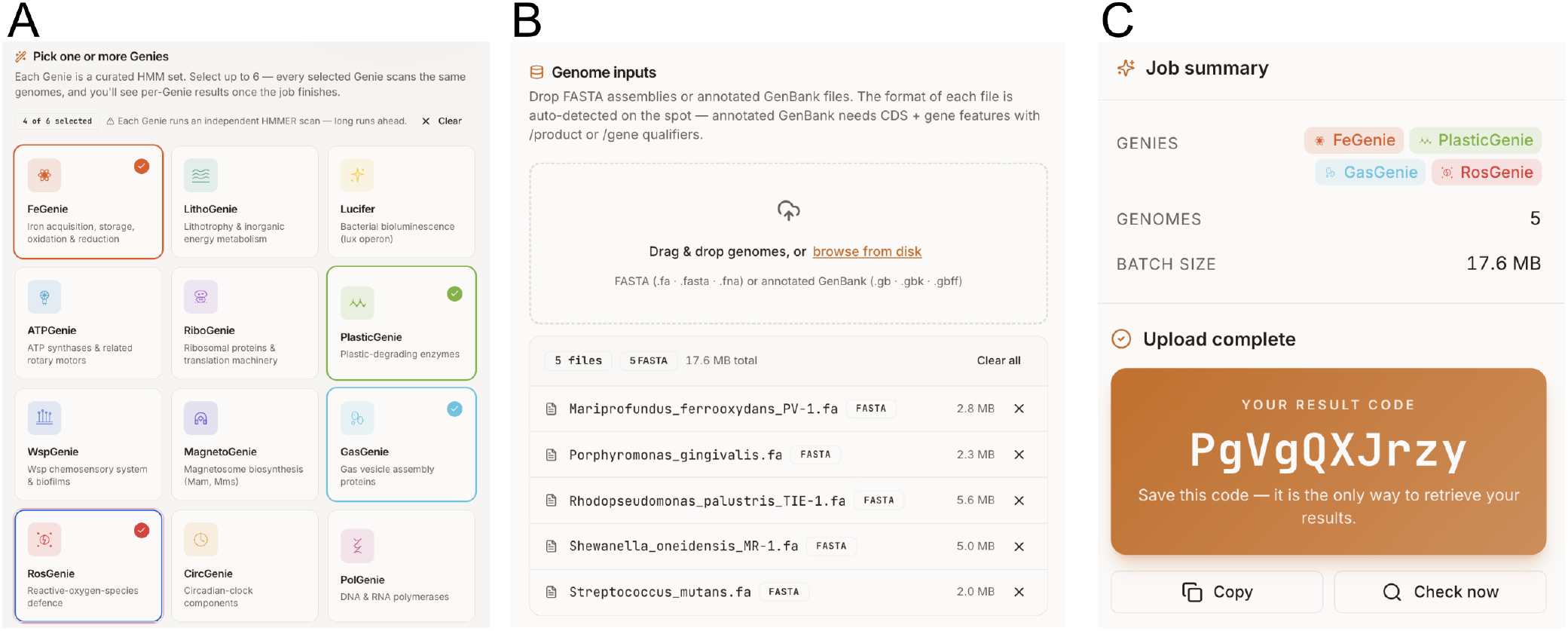
The web application interface is built on a React framework. A) Users can select up to six *genies* per analysis before (B) uploading one or more genomes (up to 100 MB of data). Data is securely uploaded to an Amazon Web Services (AWS) S3 bucket. C) Users receive a 10-digit code to access the results when the analysis is completed. This code will remain active for a week, allowing users to come back to recent analyses and save data locally.

#### Web Server Interface

In addition to the standalone command-line executable software package, MagicLamp is accessible through a web interface that supports multi-genome (up to 100 total MB) upload (**Figure 2B**). Submitted user data are processed through secure cloud infrastructure via AWS (Amazon Web Services), and results are ready for viewing immediately after analysis is complete, at an approximate rate of one bacterial genome (∼5 Mb) every 5 min. Upon submission of data, a unique results code is provided, which can later be used to retrieve results (**Figure 2C**). User data and corresponding results are not used internally and are permanently deleted after one week. Help documentation and a usage tutorial are available through the server interface.

#### Available Genie Modules

The currently available *genie* modules and their targeted functions are summarized in **Table 1**. The MagicLamp HMM library currently contains 1,075 total HMMs organized across 19 *genies* and 96 distinct subcategory assignments across all *genies* (some genies are single-category and counted as one subcategory each). The suite encompasses biological functions, including canonical microbial processes (e.g., peptidoglycan biosynthesis, motility, and sporulation), biotechnology-focused capabilities (e.g., antibiotic biosynthesis, microbial plastic degradation, and hazard resistance), and other specialized phenotypes (e.g., magnetosome formation, bioluminescence, photoheterotrophy potential, and gas vesicle regulation). Updates to existing *genies*, as well as the addition of new *genie* modules are performed on a continuing basis as new functional genes are characterized and as community needs evolve. New *genie* suggestions can be submitted directly via the web application.

## Results

For every selected *genie*, MagicLamp produces a defined set of machine-readable outputs alongside an interactive browser report. Outputs include:

1. **Per-hit annotation table (<Genie>-summary.csv)**. A long-format table in which each row represents one significant HMM hit, shown in comparison to the bit score cutoff for each HMM. Columns include the source genome or metagenome bin, ORF identifier, matched HMM, bit-score, model-specific cutoff, E-value, assigned functional category, and a proximity-cluster identifier.
2. **Aggregated gene-count matrix (<Genie>.heatmap.csv)**. A wide-format category × genome matrix that summarizes the per-hit annotation table at the functional-category level. Data within the matrix contain the number of distinct ORFs assigned to each category in each input genome, providing a compact view of function breadth, completeness (when appropriate), and abundance across datasets.
3. **Full results archive (<identifier>-results.tar.gz)**. A complete tarball (compressed archive file) of the per-run output directory, including raw HMMER tables, Prodigal-derived ORF predictions and amino-acid translations, intermediate clustering files, per-*genie* logs, and the summary CSVs.
4. **Interactive HTML report**. Each job receives a randomly generated 10-character result code that can be bookmarked, shared, and pasted back into the homepage to retrieve results. The report includes summary cards, bar charts, heatmaps, and searchable gene-call tables.

## Methods

### Aggregation of publicly available HMMs

For the majority of modules, HMMs were aggregated from public repositories — principally TIGRFAMs (Haft et al., 2003) and Pfam — after the underlying marker genes were identified through manual literature review (see Table 1 and Table S1); for each HMM file, the originating database identifier (TIGRFAM or Pfam accession) is retained in the HMM filename to preserve provenance. In the case of LithoGenie, HMMs were pulled from the metabolic-HMMs collection of Anantharaman et al. (2016). For genes for which suitable public models were unavailable, HMMs were built in-house from curated sequence sets identified through literature-based gene searches and aggregation of similar functional orthologs (see below).

### De novo construction of custom HMMs

For genes with no suitable public model (the majority of GasGenie models and a subset of PlasticGenie models), HMMs were built in-house from curated sequence sets. Reference amino acid sequences were retrieved from UniProtKB (The UniProt Consortium 2025), using reviewed (manually curated) entries. Sequence sets were expanded by a BLASTP (Altschul 1990) search against NCBI RefSeq, retaining hits with > 35% identity over > 70% of the query length.

Expanded sets were clustered with MMseqs2 (Steinegger and Söding, 2017) with > 70% amino acid identity over > 70% of the query length; alignment was performed with MAFFT, using the BLOSUM62 substitution matrix (Katoh et al., 2002), and trimmed with trimAl (Capella-Gutiérrez et al., 2009) (-gappyout), and profiles were constructed with hmmbuild (HMMER; Eddy, 2009). Each custom model was validated against the NCBI non-redundant (NR) database and against sequences from model organisms with the experimentally validated phenotype (gas vesicle production or polymer degradation). A model-specific bit-score threshold was then manually assigned to each HMM to discriminate true from false positives: we parsed NR matches to each manually constructed HMM in decreasing bit score order, and determined the optimal bit score cutoff based on where the majority of hits (sliding window of 10 HMM matches) were false positives. Existing public models that adequately captured closely related genes of interest were incorporated in the *genie* as-is, rather than rebuilt. All resulting HMMs, custom and aggregated alike, are distributed with model-specific bit-score cutoffs in the MagicLamp repository.

## Data Availability

MagicLamp is freely available at https://github.com/Arkadiy-Garber/MagicLamp. The standalone command-line version is distributed with a Conda environment containing all required software dependencies, including Python, Prodigal, and HMMER. All HMM collections currently incorporated into the toolkit are available in the repository: https://github.com/Arkadiy-Garber/MagicLamp. The web server is free and open to all users without a login requirement

## Author Contributions by genie

WspGenie was constructed by C.R. Armbruster (CAR) and A.I. Garber (AIG). FeGenie and MagnetoGenie were developed by Garber et al. (2020) and incorporated into MagicLamp by AIG. ROSGenie was constructed by AIG. Lucifer was constructed by G. Ramírez (GAR) and AIG. CircGenie was constructed by GAR and AIG. GasGenie and PlasticGenie were constructed by N. Merino (NM). LithoGenie was adapted from the metabolic-HMMs collection of Anantharaman et al. (2016) and incorporated into MagicLamp by AIG. CrisprGenie was constructed by S. Sadeghpour (SS), I.A. Viney (IAV), A. Manna (AM), M. Kreger (MK), B. Wolk (BW), E. Kim (EK), and I. Pérez-Rodríguez (IPR). SporeGenie, TaxisGenie, HazGenie, AbxGenie, and RiboGenie were constructed by IAV, AM, MK, BW, EK, and IPR. BTEXGenie was developed by Qu et al. (2026). ATPGenie, PortGenie, and RnfGenie were constructed by AIG.

## Acknowledgements

N.M. was supported by an LLNL LDRD grant (22-LW-034). This work was performed under the auspices of the U.S. Department of Energy by Lawrence Livermore National Laboratory under Contract DE-AC52-07NA27344 (LLNL-JRNL-2021449). This research was also supported by the Elliman Faculty Fellowship and start-up funds from the University of Pennsylvania awarded to I.P.-R., as well as the National Science Foundation (NSF) Award ID 2529537 to I.P.-R. C.R.A. was supported by a grant from the Richard King Mellon Foundation (Grant ID: 14323) and start-up funds from Carnegie Mellon University.

